# Development of A Decellularized Meniscus Matrix-Based Nanofibrous Scaffold for Meniscus Tissue Engineering

**DOI:** 10.1101/2020.12.23.424243

**Authors:** Boao Xia, Dong-Hwa Kim, Sonia Bansal, Yongho Bae, Robert L. Mauck, Su-Jin Heo

## Abstract

The meniscus plays a critical role in knee mechanical function but is commonly injured given its central load bearing role. In the adult, meniscus repair is limited, given the low number of endogenous cells, the density of the matrix, and the limited vascularity. Menisci are fibrocartilaginous tissues composed of a micro-/nano-fibrous extracellular matrix (ECM) and a mixture of chondrocyte-like and fibroblast-like cells. Here, we developed a fibrous scaffold system that consists of bioactive components (decellularized meniscus ECM (dME) within a poly(e-caprolactone) material) fashioned into a biomimetic morphology (via electrospinning) to support and enhance meniscus cell function and matrix production. This work supports that the incorporation of dME into synthetic nanofibers increased hydrophilicity of the scaffold, leading to enhanced meniscus cell spreading, proliferation, and fibrochondrogenic gene expression. This work identifies a new biomimetic scaffold for therapeutic strategies to substitute or replace injured meniscus tissue.

**STATEMENT OF SIGNIFICANCE:** In this study, we show that a scaffold electrospun from a combination of synthetic materials and bovine decellularized meniscus ECM provides appropriate signals and a suitable template for meniscus fibrochondrocyte spreading, proliferation, and secretion of collagen and proteoglycans. Material characterization and *in vitro* cell studies support that this new bioactive material is susceptible to enzymatic digestion and supports meniscus-like tissue formation.

## INTRODUCTION

Menisci are fibrocartilaginous tissues in the knee that transfer and redistribute load between the femur and the tibia and provide secondary stability to the joint [1]. Given these vital functions in a high load-bearing setting, menisci tears are common and occur in patients of all ages in various locations and tear patterns [2–4]. Unfortunately, the meniscus also has a limited self-healing capacity, given its dense composition and low cellularity and vascularity. Physical therapy and arthroscopic partial meniscectomy are commonly performed to alleviate symptoms [5–7]. However, these treatments do not restore the meniscus structure and function, and continued meniscus insufficiency may precipitate the onset of osteoarthritis [8, 9]. Therefore, new therapeutic strategies are needed to facilitate healing of meniscus injuries.

Over the past two decades, a number of load-bearing and/or pro-regenerative implants have emerged as commercial products to treat the injured meniscus [10]. For instance, Menaflex™ [11–14], a collagen-glycosaminoglycan (Collagen-GAG) meniscus replacement, as well as Actifit™ [11–14] and NUsurface™ [12, 14], synthetic polycaprolactone-polyurethane (PCL-PU) or polycarbonate-urethane (PCU) implants, are designed to either enhance meniscus ECM-like neo-matrix production or improve load distribution in patients who have previously been subject to partial or total meniscectomy. In addition, laboratory-based studies have developed regenerative scaffolds that utilize decellularized meniscus ECM. Examples include using the whole piece of lyophilized tissue directly as a graft [15–17], reconstituting pulverized tissue into porous or hydrogel constructs [18–20], 3D printing with ECM-based bioinks [21–25], electrospinning from solutions containing natural structural proteins similar to those present in the meniscus ECM [26–28], or a combination of the above strategies [29, 30]. Many of the studies have demonstrated improved meniscus cell or stem cell viability, infiltration, and neo-matrix deposition over time.

However, there are limitations associated with each of the above approaches. In the cases of decellularized whole meniscus transplantation, inadequate mechanical strength could lead to construct ruptures and joint deterioration, and insufficient recellularization could hamper chrondroprotective effects [15, 17]. While building bioactive scaffolds reconstituted from ECM components on hydrogel extrusion or casting platforms could provide more flexibility in terms of matching the gaps in various meniscus tears, the substrate stiffness and porosity need to be meticulously tuned to encourage cell spreading and migration [18, 20]. Therefore, it is important to devise a degradable material and fabrication method to make a scaffold that does not rupture but still supports meniscus cell activities through its bioactive open pore surface features.

Electrospinning is an advanced fabrication technique widely used to produce scaffolding materials that possess a nanofibrous structure comparable to the ECM of fibrous connective tissues. Researchers have recently spun natural materials such as gelatin, collagen, and ECM together with synthetic polymers to produce biomimetic scaffolds for repair and regeneration [26, 27, 31–34]. These scaffolds are biocompatible and show enhanced cell adhesion and proliferation compared to their purely synthetic counterparts, possibly due to enhanced hydrophilicity and bioactivity of the scaffolds. One limitation to this strategy, however, is the use of toxic organic solvents, for example, trifluoroethanol (TFE) and hexafluro-2-propanol (HFIP) in the preparation of the electrospinning solution, which poses hazards to the researchers via inhalation and may impede regulatory approval of these approaches [35–37]. Such methods could be improved by the derivation and testing of “green” solvents, and the optimization of conditions under which such solvents homogenize both natural and synthetic scaffold components.

To address this need, in this study, we developed a safe and efficient method to incorporate decellularized bovine meniscus ECM as a biomimetic component within a nanofibrous scaffold. We then developed a process to fabricate a decellularized meniscus ECM/poly(ɛ-caprolactone) (dMEP) nanofibrous scaffold for meniscus regeneration by co-electrospinning the homogenized solution. Given that the scaffold will eventually be implanted in a hydrated *in vivo* environment, we tested two common collagen crosslinkers, glutaraldehyde (GA) and genipin (GP), and evaluated their ability to maintain fiber morphology, as well as their biocompatibility with meniscus cells. We then compared these dMEP scaffolds with their PCL-only counterparts via a series of material characterization tests and *in vitro* cell studies. We hypothesize that the dMEP scaffolds would promote meniscus cell spreading, proliferation, and differentiation to a greater extent than PCL-only scaffolds. And therefore, this novel combination of bioactive content with advanced scaffold fabrication techniques may generate material frameworks that can optimally promote meniscus tissue formation and regeneration.

## MATERIALS AND METHODS

### 2.1 Tissue Decellularization and Verification

Decellularized meniscus ECM (dME) was generated using a protocol modified from Wu et al. [18]. In brief, menisci were harvested from juvenile bovine knee joints (Research 87, 2-3 months old) and were minced into cubes of approximately 1 mm^3^ (Fig. 1A i-ii). To achieve decellularization, meniscus cubes isolated from each meniscus were stirred in a 1% SDS/PBS (w/v) solution for 72 h, with the solution refreshed every 24 h. Next, the tissue was washed in a 0.1% EDTA/PBS (w/v) solution for 24 h (Fig. 1A iii). Finally, the tissue was rinsed in an excess of distilled water for 12 h and then lyophilized for 72 h. Dried tissue was ground into fine powder using a freezer-mill (SamplePrep™ 6770 Freezer/Mill™, precool time = 1 min, runtime = 2 min, rate = 14 cycles/second). To verify that menisci were appropriately decellularized, the meniscus cubes pre and post decellularization were fixed in phosphate buffered paraformaldehyde (PFA) at 4°C overnight, cleared in CitriSolv and embedded in paraffin for sectioning (thickness = 6 μm). Cell removal was confirmed by hematoxylin and eosin (H&E) staining and was quantified by counting the remaining nuclei on histological sections stained with 4,6-diamidino-2-phenylindole (DAPI) (n = 5, imaged at 10X on a Nikon Ti inverted fluorescence microscope). Preservation of collagen and proteoglycans in the decellularized tissue was determined via picrosirius red (PSR) and alcian blue (AB) staining, respectively [34, 38]. Images were captured with an Eclipse 90i upright microscope.

**Fig. 1:**
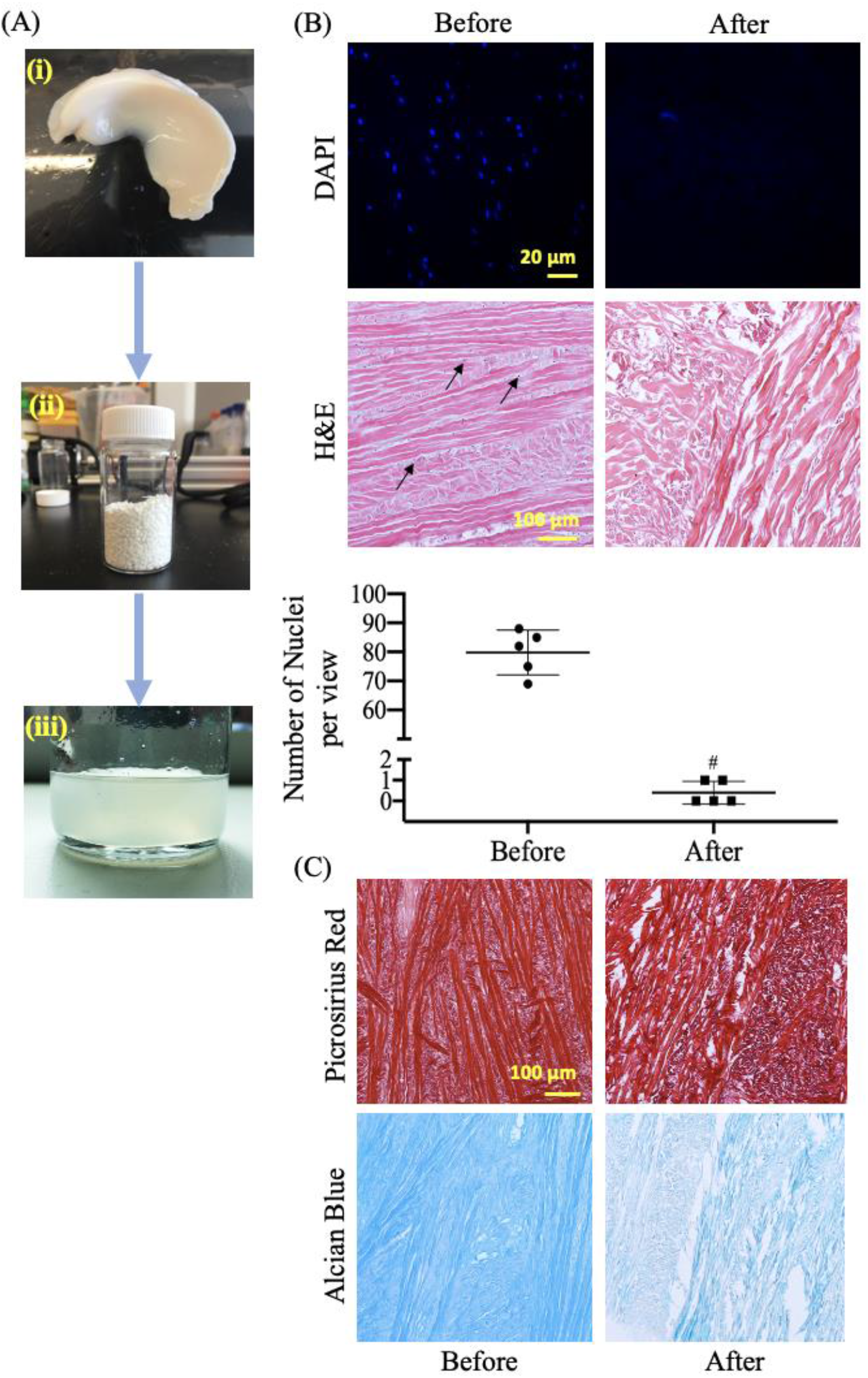
(A) Preparation of electrospinning solution: (i) whole bovine meniscus (ii) lyophilized decellularized meniscus cubes and (iii) ECM in AED solution. (B) Representative images of DAPI staining and quantification of nuclei per view [#: p<0.05 vs. before, n = 5, mean±SD, experiments were carried out at least in duplicate], and H&E staining (arrows: nuclei) before and after decellularization. (C) Representative images of picrosirius red staining for collagen and alcian blue staining for proteoglycan content before and after decellularization.

### 2.2 Electrospinning decellularized meniscus ECM – polycaprolactone (dMEP) nanofibers

Electrospinning solutions were prepared using the protocol optimized from Binulal et al. and contained both ECM and poly(ε-caprolactone) (PCL, 80kDa) [39]. Specifically, a 28% w/v dMEP (50:50) mixture was prepared by dissolving 1.4 g of dME powder in 10 mL of a diluted acidic solution (Acetic Acid : Ethyl Acetate : ddH2O (v/v/v) = 3 : 2 : 1) at 45°C for three days. Next, 1.4 g of PCL was added to the solution and stirred at the same temperature for an additional two days. Nanofibrous scaffolds were produced via electrospinning at a voltage of 15 kV, a needle-to-collector distance of 9 cm, and a flow rate of 2.2 mL/h, with randomly-organized fibers collected onto a grounded mandrel rotating at a slow speed. Relative humidity was maintained between 19% - 24%. Additional electrospun PCL-only scaffolds were spun with a similar average fiber diameter and alignment as a control. For this, 2.4 g of PCL was dissolved in 10mL of Acetic Acid/Ethyl Acetate (1:1) solution at 45°C for two days. Afterwards, this solution was electrospun at a voltage of 15 kV, a needle-to-collector distance of 14 cm and a flow rate of 2.2 mL/h. The scaffolds were removed from the mandrel and maintained in a vacuum chamber at room temperature prior to further analysis [18, 40].

### 2.3 Crosslinking and Morphological Observation of dMEP Nanofibers

Due to the instability of collagen and GAG in an aqueous environment, an effective crosslinker is required to keep the scaffolds intact [41–44]. To accomplish this, two collagen crosslinkers were tested: glutaraldehyde (GA, Sigma-Aldrich) and genipin (GP, Wako Chemicals and Challenge Bioproduct). The dMEP scaffolds were either punched into rounds (∅ = 1 cm) for surface characterization and cell studies or cut into strips (40 mm x 5 mm) for mechanical testing and histological analysis. For GA crosslinking, samples were incubated in a chamber containing 50:50 GA: dH_2_O vapor at 25°C for 48 h and quenched in 0.1 M glycine for 1 h. For GP crosslinking, samples were submersed in 0.4 M GP ethanol solution at 37°C for 48 h. Both the dMEP and control (24% PCL only) groups were rehydrated in a series of EtOH/dH_2_O solutions with graduated, sequentially decreasing concentrations (100% to 0%) for further material characterization and *in vitro* cell studies [33]. Samples were not directly matched (paired) in these studies, though all were derived from the same fabrication runs.

### 2.4 Analysis of dMEP Nanofibers with Scanning Electron Microscopy

The surface morphology of the scaffolds before and after crosslinking was examined by sputter coating the samples with 8 nm of iridium on an EMS Quorum Q150T ES sputter coater, then imaging with an FEI Quanta FEG 250 scanning electron microscope (SEM) at a distance of 10 mm and magnification of 1000x. Comparison of fiber diameters were done via manual contouring and measurement in FIJI [45].

### 2.5 In vitro Enzymatic Degradation of dMEP Nanofibers

To explore the degradation behavior of dMEP scaffolds, acellular PCL and GA or GP crosslinked dMEP scaffolds (n = 4/group) were digested in 2 mg/mL collagenase type 2 solution (Worthington) for 24 h at 37°C, and then digested in 100 ug/mL proteinase K in tris-HCl overnight at 60°C. A scaffold from each group was used for SEM imaging to visualize changes in fiber morphology and structure after enzymatic degradation, and the remaining were used for the orthohydroxyproline (OHP) assay to quantify remaining collagen [46]. These scaffolds were compared against scaffolds from each group that were not collagenase-treated through both SEM assessment and the OHP assay.

### 2.6 Assessment of Scaffold Hydrophilicity

Hydrophilicity, or surface wettability, may influence the composition of the adsorbed protein layer, which could in turn regulate how cells respond to the material [47]. Thus, contact angle analysis was performed to compare the surface wettability of the dMEP nanofibrous scaffold to the PCL-only scaffold, after rehydration and air-drying (n = 4/group). A drop of 10 μL dH2O was gently deposited onto a piece of air-dried scaffold (diameter = 1 cm) and time lapse images were taken for 90 seconds at 30 second intervals. Contact angle was measured using the angle tool in ImageJ [45].

### 2.7 Mechanical Evaluation of Scaffolds

Uniaxial tensile testing was performed on rectangular-shaped dMEP and PCL-only scaffold strips (size = 40 mm x 5 mm, n = 6-7/group). A dry (before crosslinking) scaffold group and a wet (after crosslinking) scaffold group were included in this analysis. The thickness and width of the scaffold were measured using a custom laser thickness measurement system, and the average cross-sectional area was calculated with a MATLAB code [33, 46] (shown in **S Fig. 3**). The test samples were then gripped at both ends on an Instron 5542 material testing system with a gauge length of 20 mm. Samples were extended to failure at a constant strain rate of 0.2%/sec. The elastic modulus was calculated from the linear region of the stress-strain curve [40].

### 2.8 Assessment of Cell Adhesion and Proliferation

To evaluate cell adhesion and proliferation on the scaffolds, bovine meniscus fibrochondrocytes (bMFCs) were isolated from the outer region of freshly isolated medial and lateral juvenile (2-3 months) bovine meniscus tissue and expanded to passage 2 (P2) prior to seeding [40, 48]. After rehydration and triple-rinsing in sterile PBS, the dMEP composite GA-crosslinked and GP-crosslinked, and PCL-only scaffolds were UV sterilized for 30 minutes prior to cell seeding. To asses cell adhesion and spreading, 500 bMFCs were seeded onto each patch (∅ = 1 cm), submerged in a chemically defined growth factor-free media (High glucose Dulbecco’s minimal essential medium (DMEM), 0.1mM Dexamethasome, 50ug/mL Ascorbate-2-Phosphate, 40ug/mL L-Proline, 100ug/mL Sodium Pyruvate, ITS+ Premix, 1% penicillin/streptomycin/fungizone (PSF), 1.25 mg/mL bovine serum albumin (BSA), and 5.3ug/mL linoleic acid) [38, 48], and incubated at 37°C under 5% CO_2_ for 1, 3, or 6 hours. At each time point, cells were fixed in 4% paraformaldehyde (PFA) and stained with Alexa Fluor 488 Phalloidin to visualize the cytoskeleton. To quantify cell spreading, images were captured at 20x magnification on a Leica DM 6000 widefield microscope and cell area, aspect ratio, and solidity were analyzed in FIJI using these images (n = 27-30/group/time point) [51].

Cell proliferation on the PCL-only and dMEP scaffolds was evaluated using a cell counting kit (Cell Counting Kit – 8 (CCK-8), Sigma) [33, 34]. Prior to CCK-8 assay, bMFCs were seeded onto the round patches (∅ = 1 cm, 5000 cells/patch, n = 6/group), submerged in growth factor-free media (DMEM +1% PSF +10% fetal bovine serum), and incubated at 37°C and 5% CO2 for 1 or 3 days. At the end of the incubation period, each scaffold was submerged in the CCK-8 reagent in a 96-well plate at 37°C for 2 h, and the absorbance (which is directly proportional to living cell population) was read on a Synergy H1 microplate reader at 450 nm.

To evaluate cell viability on scaffolds, 20,000 P2 bMFCs were seeded onto round scaffold patches and incubated in basal growth media for 1 or 7 days. On the day of imaging, the scaffolds were submerged in a Live/Dead staining solution (PBS : EH: Calcein-AM = 1mL : 2μL: 0.5μL, Sigma) for 45 minutes at 37°C, and imaged with a Leica DM 6000 widefield.

### 2.9 Assessment of Transcriptional Activation and Gene Expression

To assess the influence of the dMEP scaffold on transcriptional activities in cells, 500 bMFCs were seeded onto scaffolds (∅ = 1 cm) and incubated in basal growth media for 24 h. Afterwards, they were permeabilized and fixed in a freezing cold methanol/ethanol (50:50) solution for 6 minutes. Samples were then blocked with 1% BSA, followed by a triple rinse in PBS before immunofluorescence staining. Cells were stained first with transcriptional activation markers, Acetyl-H3K9 (AC-H3K9) (Invitrogen # MA5-11195, 1:400) or RNA polymerase II (POL-II) (Invitrogen # MA5-23510, 1:500) for 1 h at room temperature [50, 51]. Next, after a triple rinse, Alexa Fluor 488 goat anti-rabbit secondary antibody (Invitrogen A-11008, 1:200) or Alexa Fluor 546 goat anti-mouse secondary antibody (Invitrogen A-11030, 1:200), respectively, was added at room temperature for an additional hour [25, 26]. Images were captured at 100x magnification using a Leica DM 6000 widefield microscope and fluorescence intensity was analyzed in FIJI (n = 18-22/group) [51].

For fibrochondrogenic gene expression analysis, 5,000 bMFCs were seeded onto the scaffolds and cultured in a chemically defined media containing 10 ng/mL TGF-β3 for one week (∅ = 1 cm, n = 6/group) [33, 38, 48]. RNA was extracted from samples preserved in TRIzol™ Reagent (Invitrogen), and mRNA was quantified on an ND-100 Nanodrop Spectrometer. cDNA was synthesized using a SuperScript™ IV First-Strand Synthesis System (Invitrogen), and amplified using an Applied Biosystems Step One Plus real-time PCR system. Amplification curves for Collagen I, Collagen II, Aggrecan and CTGF were analyzed in the linear region of the amplification and normalized against the housekeeping gene glyceraldehyde-3-phosphate dehydrogenase (GAPDH) [34, 49].

### 2.10 Assessment of Matrix Content

To evaluate matrix production and dECM retention in and on the scaffolds over time, 5,000 bMFCs were seeded onto three groups of scaffolds (40 mm x 5 mm) and incubated in a chemically-defined culture media containing TGF-β3 for up to 4 weeks. At each time point, the scaffolds were removed, dried, and weighed on an analytical balance (n = 5-6/group). The scaffolds were then digested in 100ug/mL proteinase K in tris-HCL overnight at 60°C. After that, the OHP assay was performed for collagen quantification and 1,9-dimethylmethylene blue (DMMB) assay was performed for GAG quantification [46, 52]. DNA content in the digest was quantified with the Quant-iT™ PicoGreen™ dsDNA assay (Invitrogen).

### 2.11 Statistical analyses

Statistical tests were performed in the PRISM 8 software. Specific analyses included a t-test to confirm decellularization, compared scaffold mechanical strength (before crosslinking), evaluated chondrogenic gene expression. For other outcomes with multiple groups, a one-way ANOVA was used to compare fiber diameter, OHP content following collagenase treatment, scaffold mechanical strength (after crosslinking), and transcriptional activation. For other outcomes, a two-way ANOVA was used to assess differences in hydrophilicity, cell adhesion, long term matrix content. Either Tukey’s or Kruskal-Wallis post hoc comparisons were used with a confidence interval of 95%.

## RESULTS

### 3.1 Characterization of Decellularized ECM

After the decellularization process, cell removal in the meniscus ECM was first confirmed by counting the remaining DAPI stained nuclei or visualizing nuclei by H&E staining of histological sections. While the freshly harvested juvenile meniscus tissue contained a large number of fibrochondrocyte-like cells (MFCs), DAPI staining showed that the cells were effectively removed from the tissue with the decellularization process, with 0.5% of the nuclei remaining in five biological replicate groups. Effective cell removal was also confirmed by H&E staining (**Fig. 1B**). The collagen content and architecture were preserved after the decellularization treatment (**Fig. 1C, top row**). In contrast, there was a noticeable decrease in alcian blue staining intensity, indicating a loss of proteoglycan content with decellularization (**Fig. 1C, bottom row**).

### 3.2 Electrospun dMEP Nanofibers

To fabricate nanofibrous scaffolds containing decellularized native meniscus ECM, the dMEP mixture was prepared and electrospun (**Fig. 2A**). Suitable formulations were selected based on the ability to spin a scaffold without interruption of fiber formation, appropriate fiber diameter in the collected scaffold, and the lack of irregularities/inclusions in the formed mat, which are indicative of an unstable Taylor cone.

**Fig. 2:**
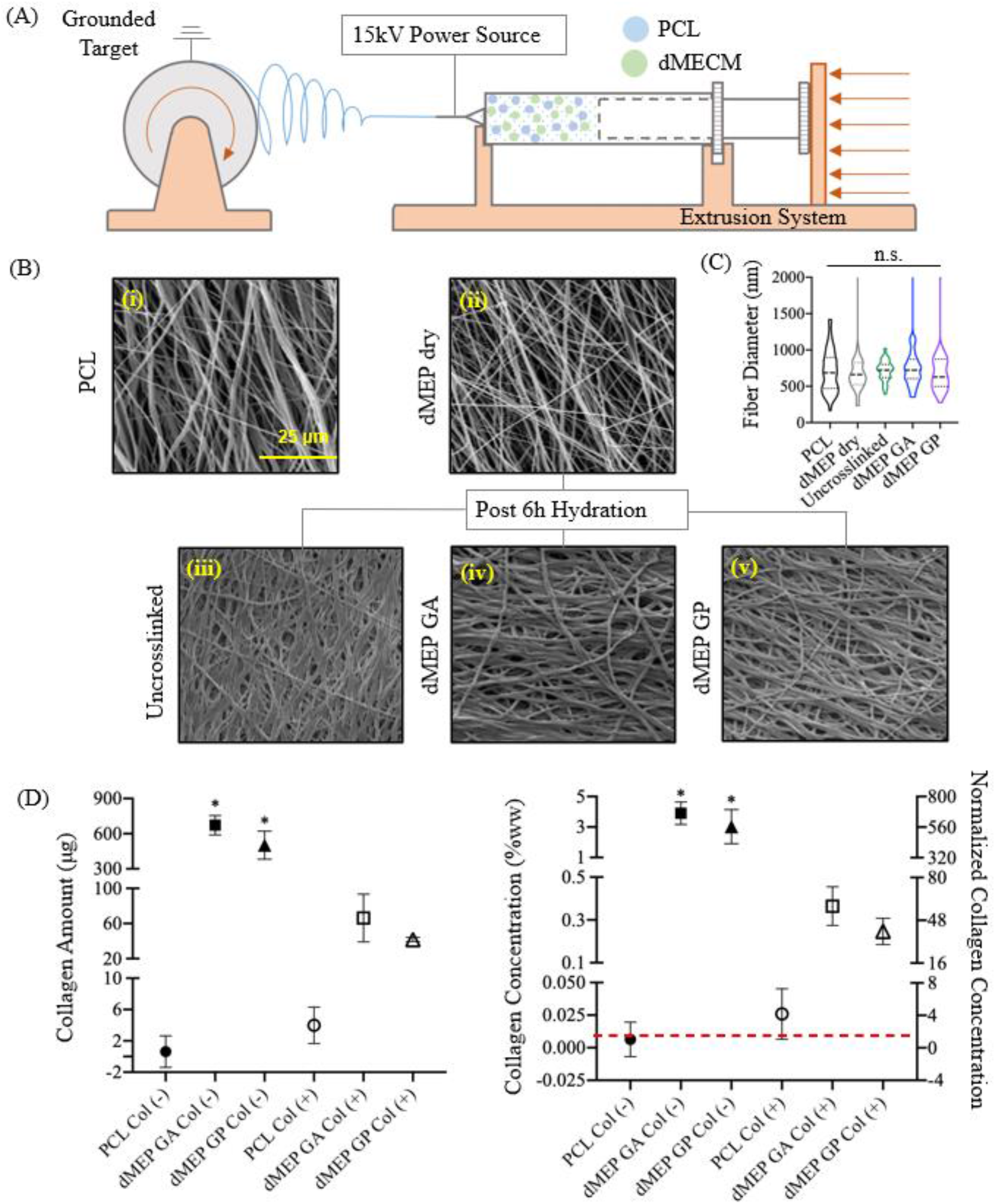
(A) Diagram of electrospinning of the dMEP mixture; (B) Representative SEM images of (i) 24% PCL control fibers, (ii) uncrosslinked, dry dMEP fibers, iii) uncrosslinked dMEP, (iv) Glutaraldehyde (GA) crosslinked dMEP fibers, and (v) Genipin (GP) crosslinked dMEP fibers post 6h submersion in water; (C) Comparison of PCL and dMEP fiber diameter [n = 50, dashed line marks median, dotted lines mark 25 percentile and 75 percentile]; (D) Collagen amount and concentration of acellular PCL and dMEP scaffolds before and after 24h of 2mg/mL collagenase type 2 treatment [right Y axis normalization to week 1 PCL group, *: p < 0.05, vs. PCL, n = 4 per group, mean±SD].

### 3.3 Assessment of Fiber morphology via SEM

The nanostructure of the electrospun scaffolds pre- and post-crosslinking was examined by scanning electron microscopy (SEM). Fiber diameter and orientation of the uncrosslinked and dry dMEP scaffold were similar (p>0.05) to those of the PCL control scaffold (**Fig. 2B i-ii, 2C**). SEM images demonstrated that crosslinking with either GA or GP preserved the fibrous morphology, while hydration of uncrosslinked dMEP scaffolds resulted in substantial changes in fibrous morphology (**Fig. 2B iii-v**). There was minimal batch to batch variability in terms of fiber diameter for dMEP scaffolds electrospun on different dates under the same conditions (**S Fig. 1**)

### 3.4 Assessment of Collagen Content in dMEP Nanofibers

The initial collagen content of dMEP nanofibers was considerably higher than PCL nanofibers. This was verified by collagenase treatment of the dMEP fibers, where the collagen content decreased 10-fold after collagenase treatment (p<0.05) [Col(+), **Fig. 2D**]. This finding also suggests that this biomimetic component of the dMEP fibers is accessible and may be degraded by collagenase secreted by endogenous meniscus cells or cells seeded onto the scaffold. dMEP nanofibers were straighter after collagenase treatment, with a decrease in fiber diameter, suggesting there might have been some amount of pre-tension within fibers as a result of the electrospinning and crosslinking process (**S Fig. 2**).

### 3.5 Assessment of Scaffold Hydrophilicity

The hydrophilicity of a scaffold is an important factor for cell attachment, spreading, and proliferation [53] and also affects oxygen and nutrient transfer within the scaffold [54]. The wettability test showed a decrease of the contact angle over time for all three groups. Water was absorbed into the dMEP scaffold more rapidly (p<0.05) than the PCL scaffold (**Fig. 3A**), indicating that the addition of dECM improved hydrophilicity. Of note, the initial contact angle for the PCL scaffolds were higher than for the dMEP scaffold with either GA or GP crosslinking (p<0.05, **Fig. 3B**), and this relationship was conserved for final water contact angles measured at 90 seconds.

**Fig. 3:**
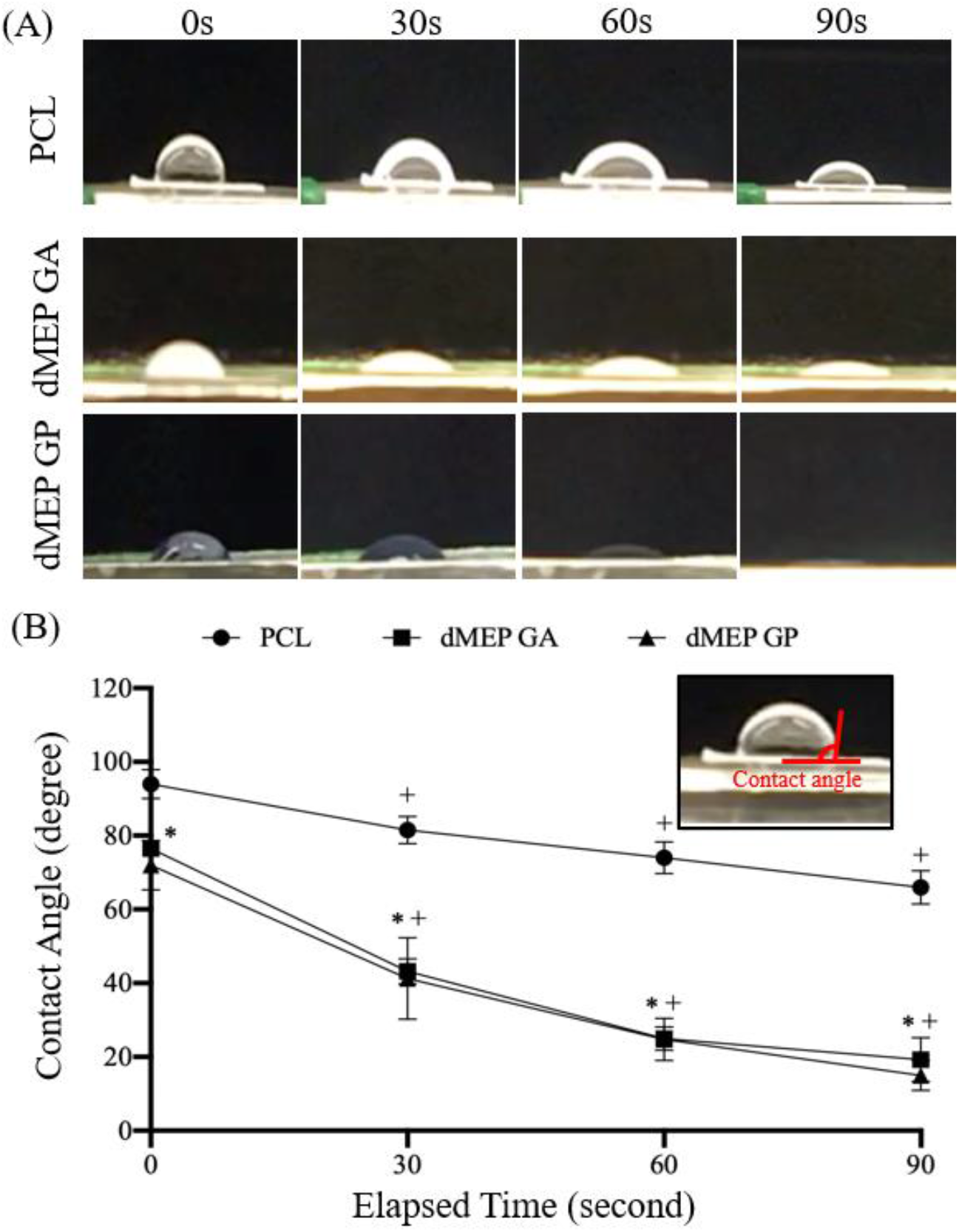
(A) Representative images of contact angle change on the water drop on 3 groups of scaffolds (PCL control, GA- or GP-crosslinked dMEP) at elapsed time points 0, 30s, 60s, and 90s. (B) Quantification of contact angle changes with time [*: p < 0.05 vs. PCL, +: p < 0.05, vs. 0s, n = 4 per group, mean±SD].

### 3.6 Mechanical Strength of Scaffolds

Additionally, crosslinking and rehydration process altered the mechanical properties of dMEP scaffolds (**Fig 4A,** individual curves in **S Fig. 3A**), in particular, ductility of dMEP scaffold was enhanced by the crosslinking process. The average tensile modulus (p<0.05) and ultimate tensile strength (p=0.09) of the dry of uncrosslinked dMEP scaffolds were higher than those of the PCL scaffolds (**Fig. 4B**). After crosslinking and rehydration, however, the modulus and ultimate tensile strength of the wet PCL scaffold were 2-3 times higher than either crosslinked dMEP scaffold (p<0.05), whose moduli were similar to one another (~0.5 MPa, **Fig. 4C**).

**Fig. 4:**
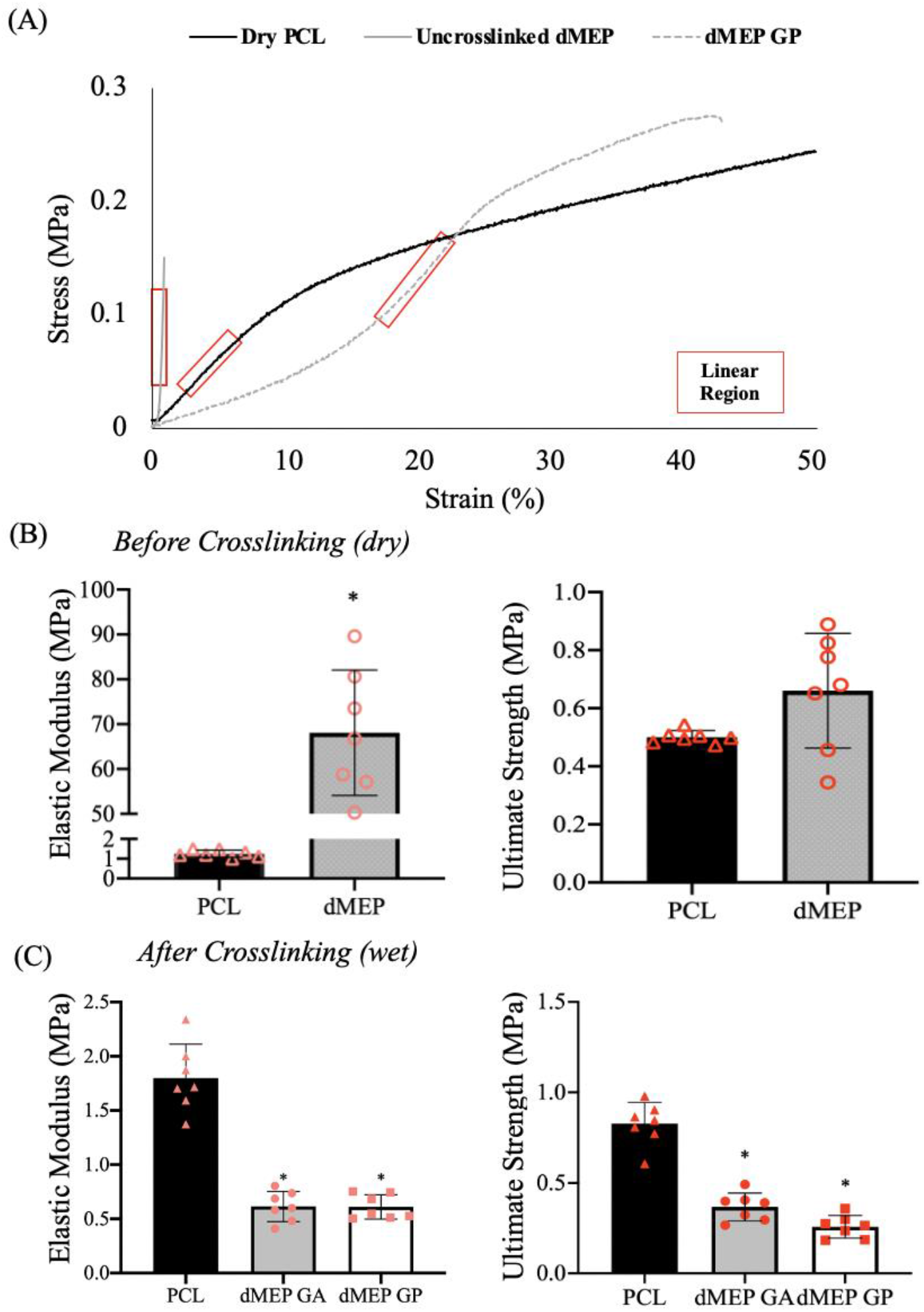
(A) Representative stress-strain curves of a PCL scaffold, an uncrosslinked dMEP scaffold and a GP crosslinked dMEP scaffold with linear regions labeled. The elastic modulus and ultimate tensile strength of PCL and dMEP scaffolds (B) before and (C) after crosslinking. [*: p < 0.05, vs. PCL, n = 6-7 per group, mean±SD, experiments were carried out at least in duplicate].

### 3.7 Assessment of Cell Spreading and Proliferation

Actin staining showed the meniscus cells attached to all 3 scaffolds over 6 hours of incubation in chemically defined serum free media. While bMFCs spread similarly on all three scaffolds, they spread more on the dMEP scaffolds, according to increased cell area (**Fig. 5A, B**, p<0.05). The cell aspect ratios were also slightly lower for dMEP scaffolds at all time points, and those cells seeded on dMEP scaffolds elongated faster (**Fig. 5A, B)**. Moreover, bMFCs proliferated faster on dMEP scaffolds over the course of 3 days (p<0.05) (**Fig. 5C**). Taken together, these data suggest the dMEP scaffolds improve cell spreading and proliferation compared to PCL-only scaffolds.

**Fig. 5:**
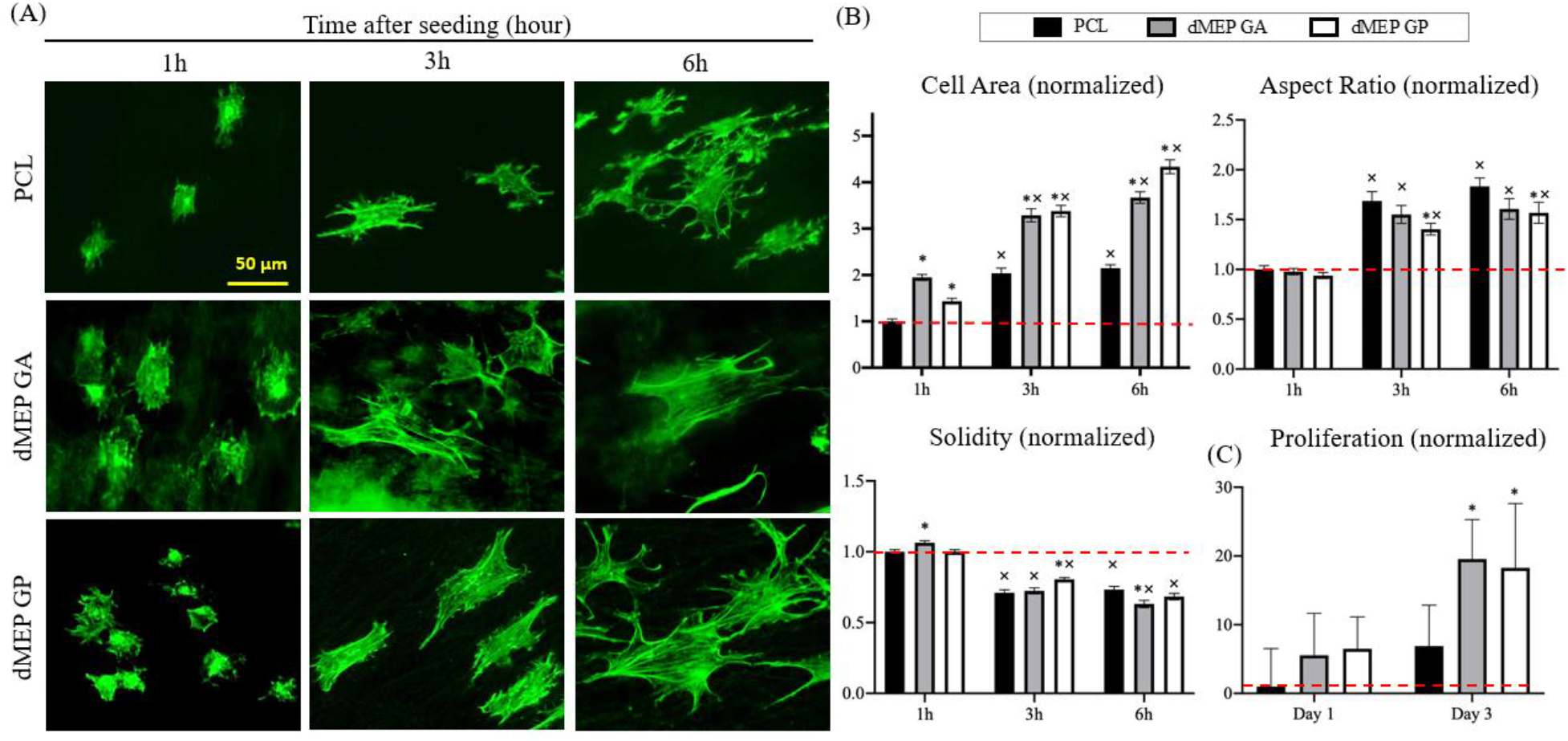
(A) Representative images of actin staining with Alexa Fluor 488 Phalloidin of bMFCs seeded onto 3 groups of scaffolds, incubated in a chemically-defined growth factor-free media for 1, 3, or 6h. (B) Quantitation of cell area, aspect ratio and solidity of bMFCs seed onto the scaffolds and incubated for 1, 3, or 6h. All normalized to the 1h PCL group [*: p < 0.05, vs. PCL; x: p < 0.05, vs. 1h, n = 27-30 per group, mean±SEM]. (C) Absorbance reading at 450 nm from a CCK-8 assay terminated at day 1 or 3 for bMFCs incubated in the growth factor-free media. Normalized to the Day 1 PCL group. [n = 6 per group, mean±SD]

### 3.8 Assessment of Transcriptional Activation and Gene Expression

Compared to the PCL scaffolds, the fluorescence intensity of both AC-H3K9 or POL-II transcriptional activation makers [50] was higher in the bMFCs cell nuclei seeded on dMEP scaffolds (p<0.05) (**Fig. 6A, B**). Further, the expression of type-I collagen (Col I), type-II collagen (Col II), aggrecan (AGC) and connective tissue growth factor (CTGF) were all ~20% higher in cells on dMEP scaffolds compared to cells on PCL control scaffolds at 1 week (**Fig. 6C**, p<0.05). Live and dead staining showed that the viability of cells was high 24 h post seeding and was further enhanced by the dMEP scaffold crosslinked by GP (**S Fig. 4**).

**Fig. 6:**
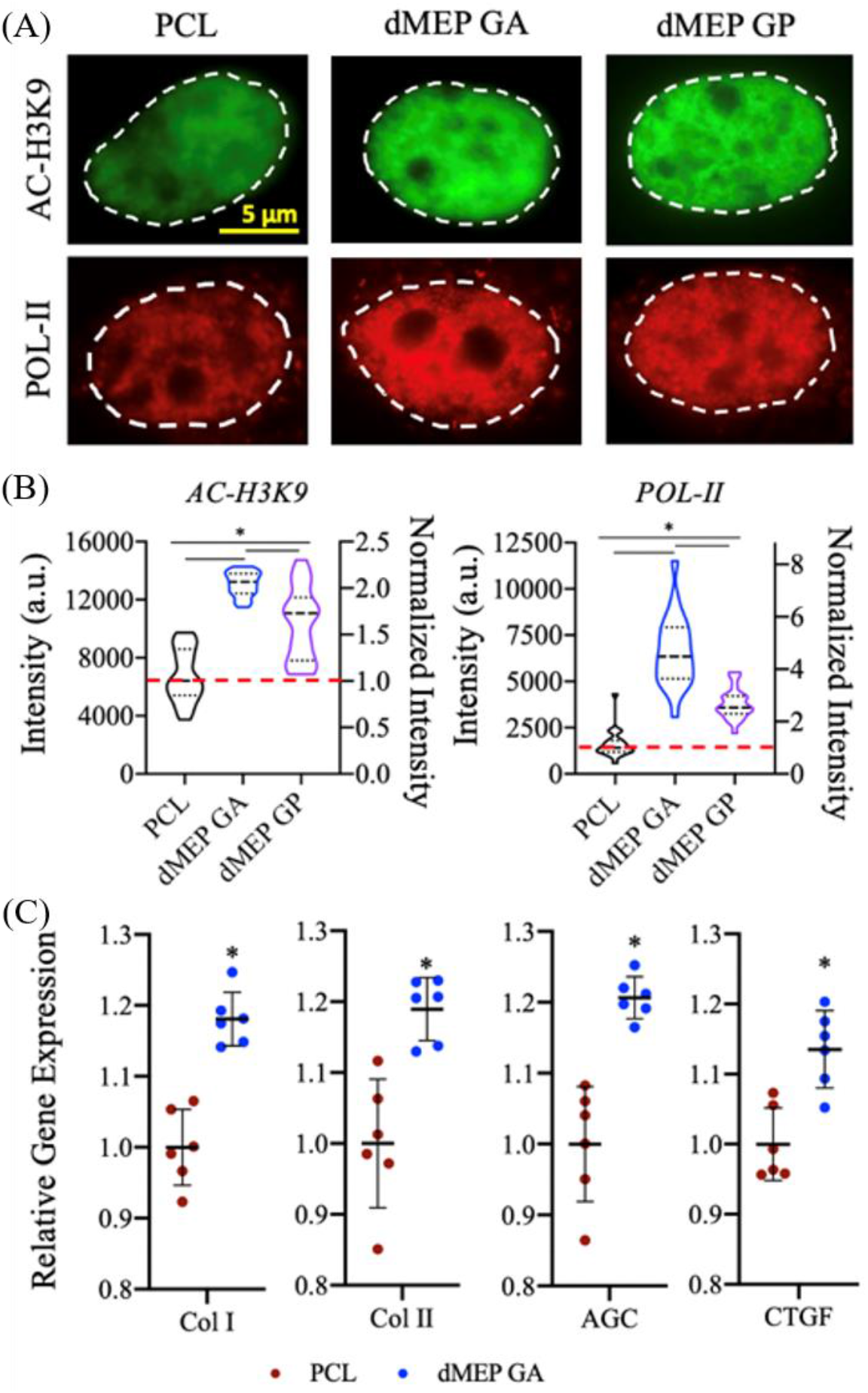
Representative (A) images and (B) quantification of AC-H3K9 mean intensity and POL-II mean intensity of the nuclei of bMFCs cultured onto PCL, dMEP GA and dMEP GP scaffolds and incubated in growth factor-free media for 24h [*: p < 0.05, vs. PCL, n = 18-22 per group, mean±SEM]. (C) Level of chondrogenesis-related gene expression relative to GAPDH in bMFCs cultured in growth factor-containing media at day 7, normalized to PCL group. [*: p < 0.05, n = 6 per group, mean±SD].

### 3.9 Matrix Deposition with Long term Culture

MFCs seeded onto all scaffold groups proliferated steadily in growth factor-containing media, with only minor difference in proliferation rate over the course of 4 weeks (**S Fig. 5**). Total collagen content was higher in bMFC-seeded dMEP scaffolds compared to PCL-only, though the magnitude of this differences decreased over time (**Fig. 7A**). The same trend was noted after normalization by sample weight (**Fig. 7B**). Since the collagen concentration in the dMEP scaffolds was initially quite high, this may suggest that the collagen within the scaffold was being broken down by the cells during culture at a rate higher than the secretion and accumulation of new collagen, leading to a decreasing total amount. Total GAG content in PCL/ECM cell-seeded scaffolds increased slightly over 4 weeks, with a similar trend observed post normalization by sample weight (**Fig. 7C-D**). Considering that the initial GAG content in the scaffolds was very low, it is likely that the majority of GAG detected in this assay was new matrix produced by cells during the incubation phase. Noticeability, collagen amount was stable in acellular dMEP scaffolds **(S Fig. 6)**, indicating that the majority of collagen detected in the OHP assay came from the scaffold, instead of being newly secreted by seeded cells.

**Fig. 7:**
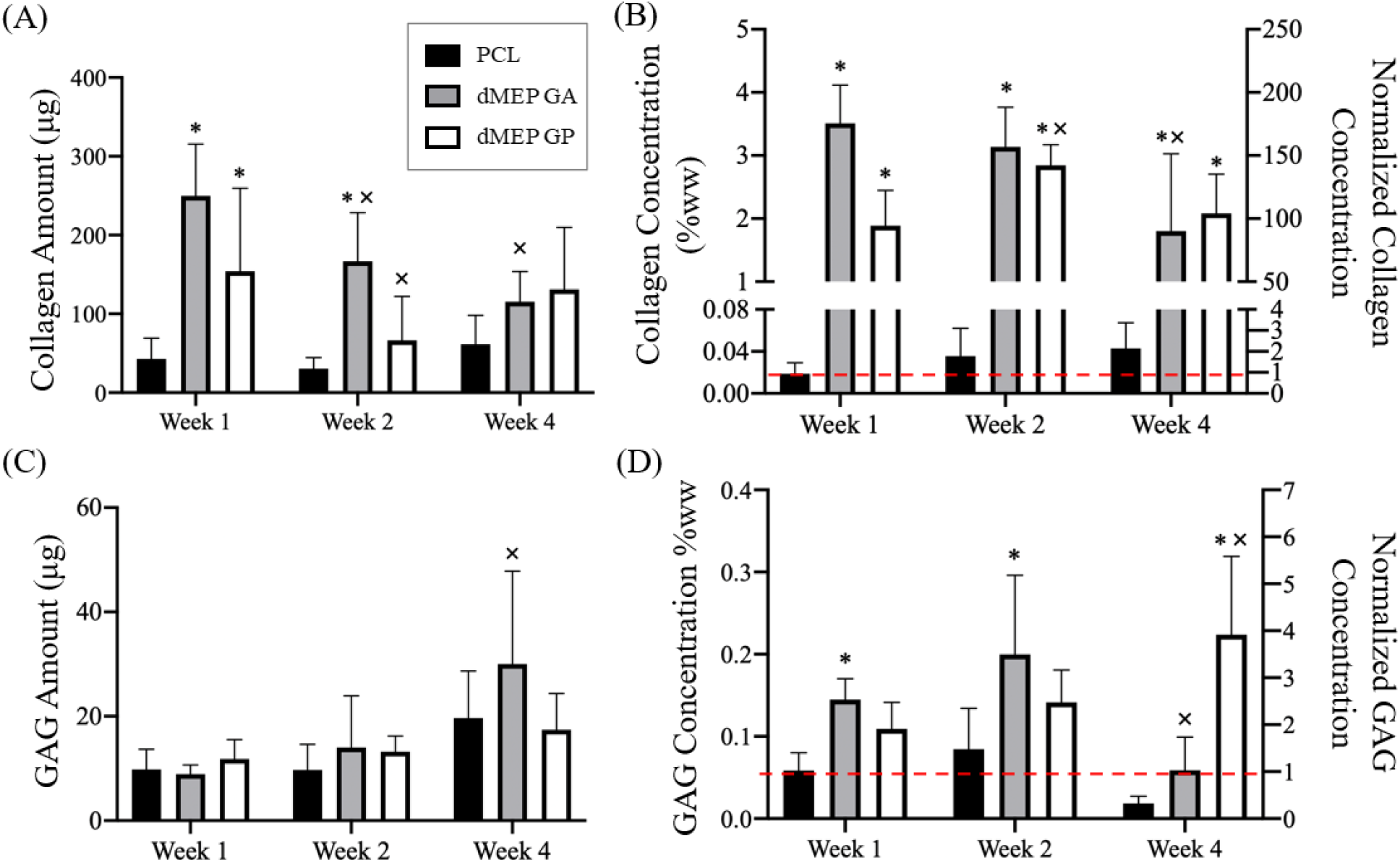
Quantification of collagen (A) amount and (B) concentration (in %wet weight, right Y axis normalization to week 1 PCL group) via OHP assay and GAG (C) amount and (D) concentration (in %wet weight, and normalized to Week 1 PCL group) via DMMB assay of bMFC-seeded scaffolds incubated in growth factor-contained media for 1, 2 and 4 weeks. [*: p < 0.05, vs. PCL; x: p < 0.05, vs. 1 week, n = 5-6 per group, mean±SD].

## DISCUSSION

In this study, we established a protocol to successfully extract decellularized meniscus ECM (dME) content. We then combined this material with a synthetic polymer (PCL) to fabricate a nanofibrous scaffold for meniscus repair. PCL was chosen among a library of biodegradable polymers for its distinct material properties; it has can be deformed elastically through physiological levels, while at the same time being degradable via enzymatic or hydrolytic mechanisms [55]. By electrospinning the (dME) material combined with the synthetic polymer from a single jet and then subsequently crosslinking the thick mat, we generated a native dME-containing nanofibrous scaffold with a similar fibrous structure to the native meniscus. Compared to the pure polymer construct, the dMEP nanofibers were more hydrophilic and bioactive, promoting cell attachment and spreading at early time points. This finding supports our hypothesis that the inclusion of dME content enhances substrate hydrophilicity (and may regulate protein adsorption) to guide initial cell attachment [56–57]. This finding is consistent with previous studies that created dME based scaffolds for musculoskeletal tissue repair (e.g. bone, muscle, tendon), in which evidence of dME promoting recellularization and organic molecule adsorption had been reported [57–59]. We also confirmed that dMEP scaffolds contained a significantly higher initial collagen components than the PCL scaffolds, and that this component was accessible to exogenous proteases. This indicates that the hybrid synthetic-biomimetic fibers can be acted on and digested by cell-produced collagenase.

With these early observations in hand, we next proceeded to longer-term evaluation. These studies confirmed that the collagen and GAG content not only started higher, but remained elevated in dMEP scaffolds over time, compared to synthetic PCL-only scaffolds. This indicates that the crosslinking utilized was effective at maintaining the dME content with culture, and that seeded meniscus cells secreted additional matrix over time. GA and GP had comparable effects in preserving dMEP fibrous morphology and strength, retaining collagen content, modifying surface wettability, and promoting cell expansion and proliferation. However, the GA group exhibited higher transcriptional activation of AC-H3K9 and POL-II than the GP group. This observation is consistent with previous studies in which the impact of crosslinkers on tissue mechanics and cellular activities were examined [33, 60]. Meniscus cells also proliferated to a greater extent on dMEP scaffolds and showed higher viability compared to the synthetic PCL-only scaffold, confirming the biocompatibility of the scaffold. The inclusion of dECM also enhanced both transcriptional activation (determined by epigenetic markers) and fibrochondrogenic gene expression (determined by RT-PCR) of meniscus cells. After 1 day, meniscus cells showed increased marks for RNA transcription overall, and by one week, showed higher levels of mRNA for collagen I, II, CTGF, and aggrecan. These data indicate that dME inclusion in the dMEP scaffold promotes initial cell activity, and that this translates into an enhanced meniscus-specific gene expression profile at one week However, these assessments of transcriptional activation and gene expression will need to be compared to expression by cells in the native meniscus, and should be expanded to include ECM degrading enzymes [60] and/or proinflammatory cytokines [61].

This study established the potential of dME inclusion to create biohybrid scaffolds which work to promote meniscus cell phenotype over long term culture and provides a foundation for further refinement and translation. One limitation of the scaffold is its physical properties: it is a sheet-based scaffold with an elastic modulus that is ~3.5% of the elastic modulus of the radial region of the juvenile meniscus, and the ultimate strength of the scaffold is ~20% of native [63]. Therefore, the dMEP scaffold would be most useful in situations where structural support is less necessary or after tissue deposition and scaffold maturation has occurred in vitro prior to implantation [64]. However, since we only explored one material fabrication technique (electrospinning), these results may be further integrated into technologies for generating meniscus-shaped constructs. For example, these fibers might be woven into thicker and aligned microfibers to further mimic the hierarchical structure of the native tissue and to reinforce its overall and directional mechanical properties [65–68]. Another important feature of the scaffolds was shown in the collagenase treatment experiment, where the dMEP fibers were thinner post-collagenase treatment (perhaps as a result of the collagen content being digested). Remodeling capacity is an important feature in any scaffold, and here cell-generated proteases may act on the dMEP scaffold to generate more space for cell infiltration [20]. To better characterize these remodeling dynamics, future *ex vivo* studies will assess the stability of the dMEP scaffold over a longer term. Importantly, however, since the remaining PCL component would be unaffected by digestion, it could continue to provide structural support during regeneration. Another feature that may be considered in the future is the heterogeneous nature of the meniscus itself, where there are inner and outer zones with distinct cell phenotypes and ECM composition [69–70]. This dMEP scaffolding system could be further refined to include zonal dME with various protein compositions to provide meniscus zone specific attributes and biological cues to infiltrating cells. Finally, through our longer-term evaluations, we confirmed that fibrocartilaginous matrix concentrations in the dMEP scaffold were high and stable during the culture periods, and that seeded cells produced new matrix. However, the balance between retention and production of neo-matrix by meniscus cells will need to be further evaluated *in vivo* to examine the performance of this scaffold in a physiologically relevant environment. Taken together, this novel combination of bioactive content with advanced scaffold fabrication generated a new translational material that can be further optimized to improve meniscus matrix formation and functional regeneration.

## ACKNOWLEDGEMENTS

This work was supported by the National Institutes of Health (R01 AR056624, K01 AR077087, R21AR077700), the Penn Center for Musculoskeletal Disorders (P30 AR069619), and the Department of Veterans Affairs (IK6 RX003416). The authors would like to acknowledge Dr. Jay Patel for discussions on experimental design, the CDB microscopy core for assistance with microscopy, Emilie Rabut for help with the freezer-mill, and Cheryl Wecksler for input on scientific language.

