## Supplementary Information for "Development of A Decellularized Meniscus Matrix-Based Nanofibrous Scaffold for Meniscus Tissue Engineering"

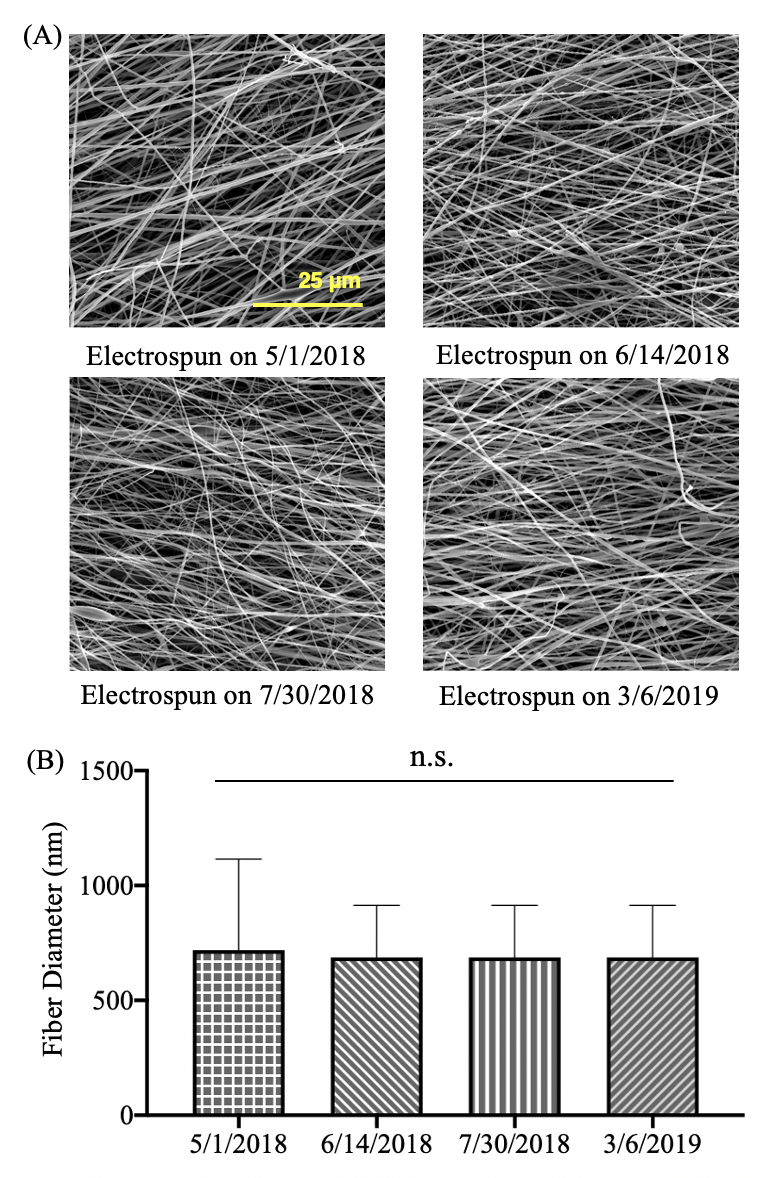

**S Fig. 1**: (A) Representative SEM images and (B) comparison of fiber diameter of dMEP scaffold electrospun on four different dates [n = 50 per group, mean+SD].

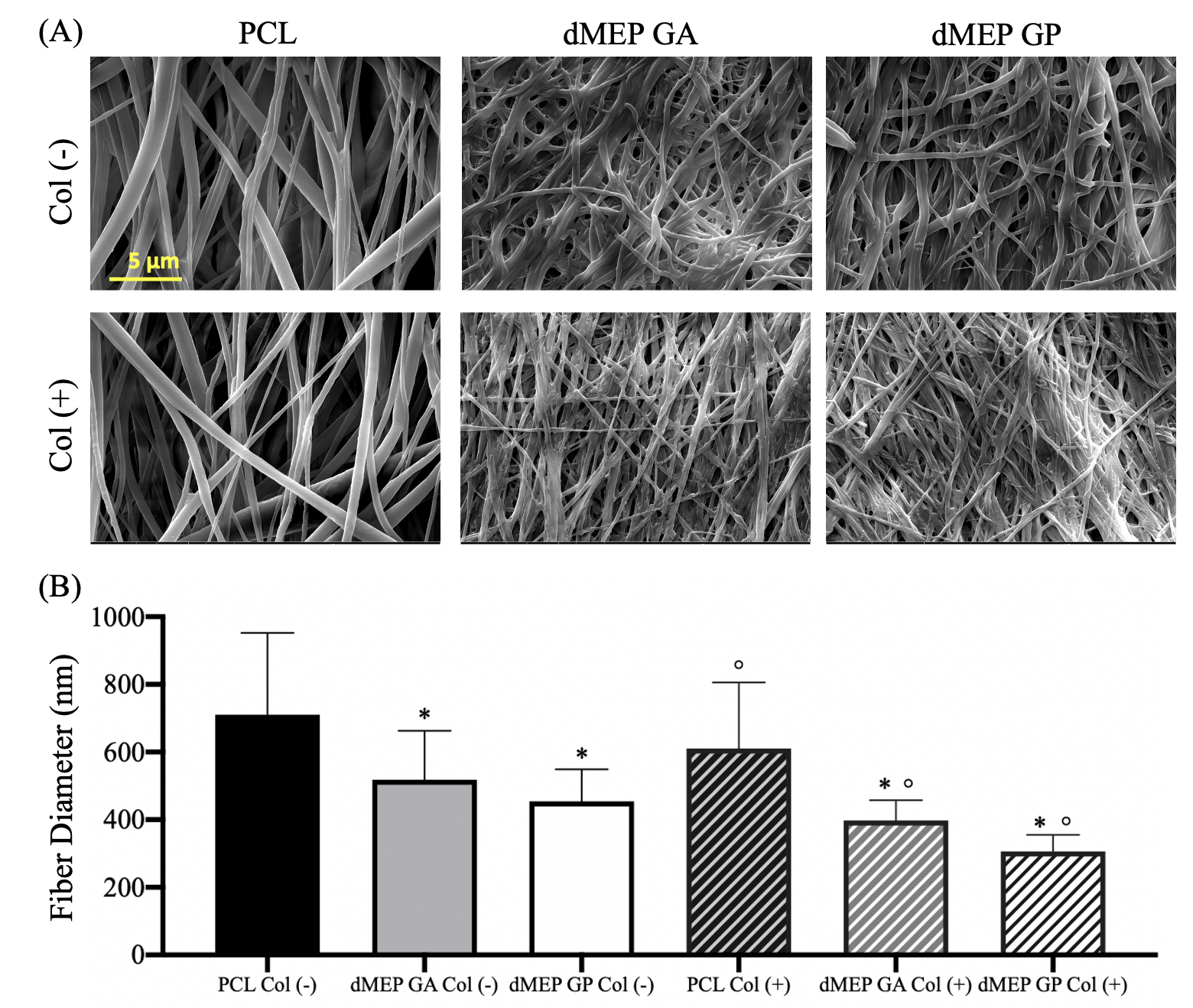

**S Fig. 2**: (A) Representative SEM images showing morphology of PCL and crosslinked dMEP nanofibrous scaffolds with/without collagenase treatment [Col (-/+)]. (B) Quantitation fiber diameter change using the SEM images above. [*: p < 0.05, vs. PCL, °: p < 0.05, vs Col (-), n = 40, mean±SD].

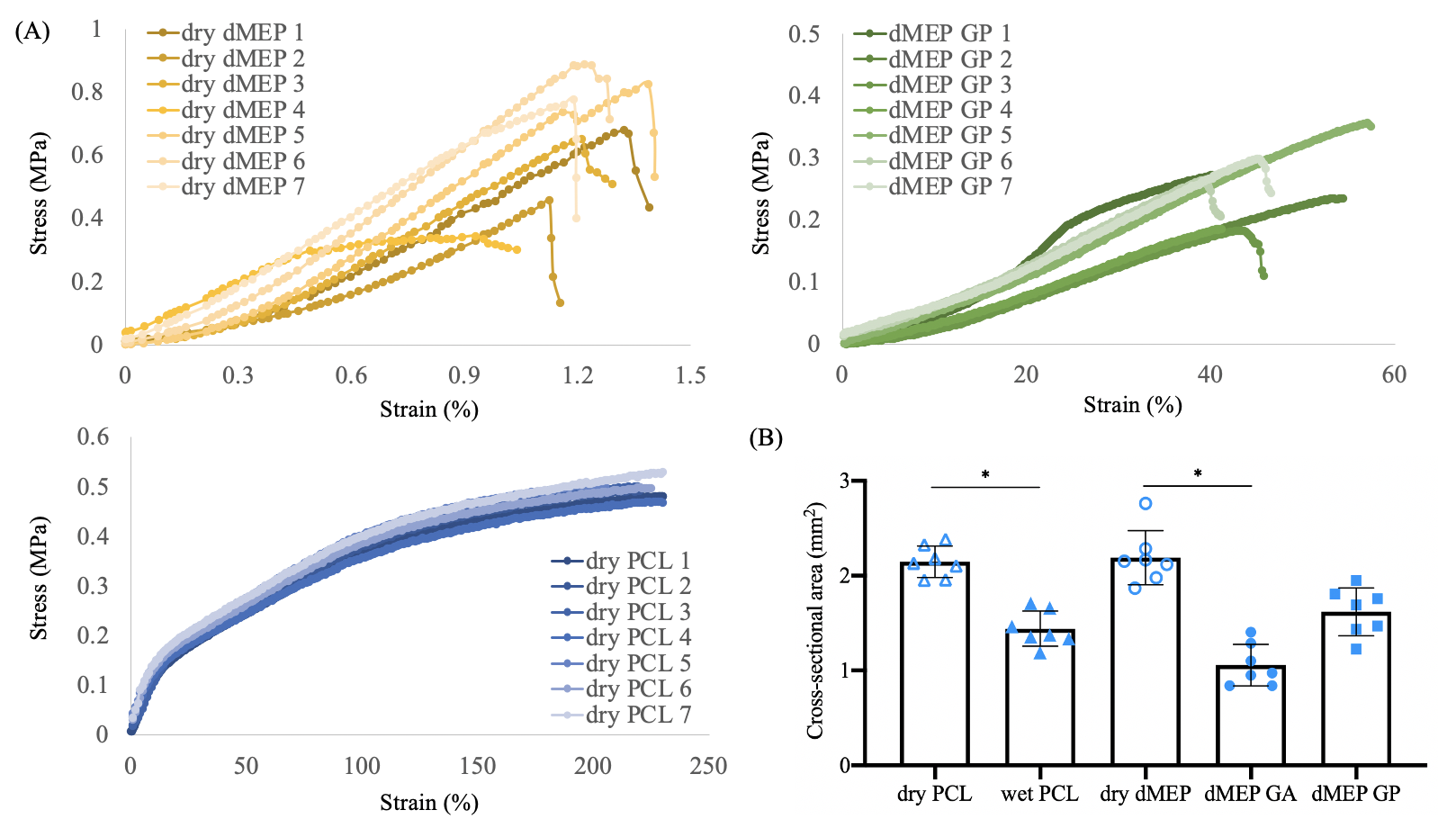

**S Fig. 3**: (A) stress-strain curves of individual uncrosslinked (dry) dMEP scaffold, GP crosslinked dMEP scaffold and dry PCL scaffold. (B) Cross-sectional area of each scaffold [*: p < 0.05, biological replicate n = 7 per group, mean±SD].

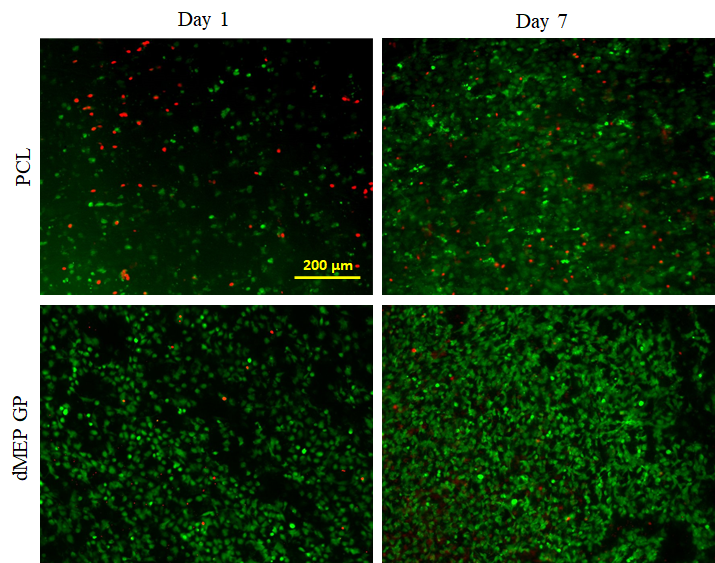

**S Fig. 4**: Live (green) and dead (red) staining of bMFCs seeded onto PCL and dMEP GP scaffolds and incubated in basal media for 1 and 7 days.

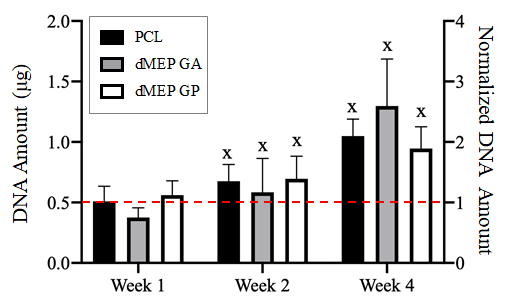

**S Fig. 5**: Quantification of DNA content (right Y axis normalization to the week 1 PCL group, red dashed line) [x: p < 0.05, vs. 1 week, n = 5-6 per group, mean ± SD].

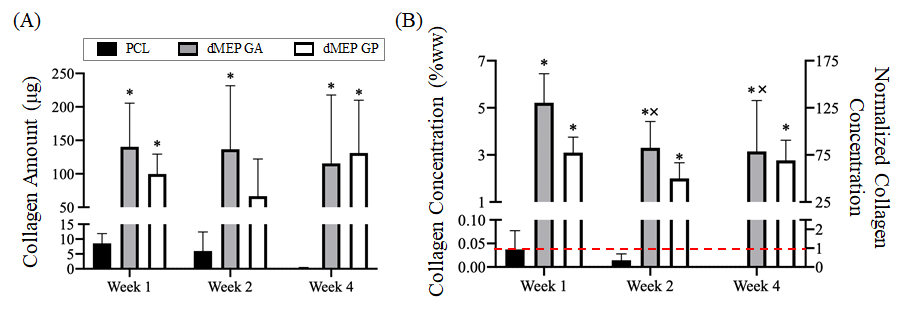

**S Fig. 6**: Collagen (A) amount and (B) concentration of acellular scaffolds (right Y axis in B represents normalization to the week 1 PCL group, red dashed line) after 1, 2, or 4 weeks of incubation in TGF-β3 containing media [*: p < 0.05, vs. PCL; x: p < 0.05, vs. 1 week, n = 6 per group, samples with undetectable collagen concentration were assigned a value of 0, mean ± SD].
